# Endoplasmic reticulum stress delays choroid development in the *HCAR1* knock-out mouse

**DOI:** 10.1101/2024.01.12.575419

**Authors:** Monir Modaresinejad, Xiaojuan Yang, Mohammad Ali Mohammad Nezhady, Tang Zhu, Emmanuel Bajon, Xin Hou, Houda Tahiri, Pierre Hardy, José Carlos Rivera, Pierre Lachapelle, Sylvain Chemtob

**Affiliations:** Program in Biomedical Science, Faculty of Medicine, Université de Montréal, Montreal, QC, Canada; School of Optometry, Université de Montréal, QC, Canada; Department of Pediatrics, Ophthalmology and Pharmacology, Centre de Recherche du CHU Sainte-Justine, Montréal, QC, Canada; Departments of Ophthalmology and Neurology-Neurosurgery, Research Institute of the McGill University Health Centre-Montreal Children’s Hospital, Montreal, QC, Canada; Program in Molecular Biology, Faculty of Medicine, Université de Montréal, Montreal, QC, Canada

## Abstract

The sub-retina, composed of the choroid and the retinal pigment epithelium (RPE), bears a critical role in proper vision. In addition to phagocytosis of photoreceptor debris, the RPE shuttles oxygen and nutrients to the neuroretina. For their own energy production, RPE cells mainly rely on lactate, a major by-product of glycolysis. Lactate in turn is believed to convey most of its biological effects via the HCAR1 receptor. Here, we show that the lactate-specific receptor, HCAR1, is exclusively expressed in the RPE cells and that *Hcar1*^−/−^ mice exhibit a substantially thinner choroid vasculature during development. Notably, the angiogenic properties of lactate on the choroid are impacted by the absence of *Hcar1*. *Hcar1*-deficient mice exhibit elevated endoplasmic reticulum (ER) stress along with eIF2α phosphorylation, a significant decrease in the global protein translation rate, and a lower proliferation rate of choroidal vasculature. Strikingly, inhibition of the Integrated Stress Response using an inhibitor of eIF2α phosphorylation (ISRIB) restores protein translation and rescues choroidal thinning. These results provide evidence that lactate signalling via HCAR1 is important for choroidal development/angiogenesis and highlight the importance of this receptor in establishing mature vision.

## Introduction

The choroid is a highly vascularized structure of the sub-retina (Nickla and Wallman 2010), which supplies O_2_ and nutrients to the Retinal Pigment Epithelium (RPE) and photoreceptors. It is primarily a vascular tissue overlaying the RPE and outer retina contributing to ocular growth and development (Zhang et al. 2022). The underlying RPE is a monolayer of post-mitotic cells that assists in the critical recycling of photoreceptor outer segments, acts as a barrier to control the passage of fluids and nutrients to photoreceptors, maintains the integrity of the choroidal vasculature, and protects the retina (Boulton and Dayhaw-Barker 2001; Strauss 2005). In this context, RPE cells also have a role in the secretion of a variety of growth and immunosuppressive factors to maintain choriocapillaris well-being (Strauss 2005; Simó et al. 2010).

The retina is the most oxygen-consuming tissue in the body and hence, metabolic disturbances are considered key contributors to ocular pathology (Flammer et al. 2013). Indeed, photoreceptor activity requires a high metabolic activity (Akhtar-Schäfer et al. 2018). RPE cells phagocytose photoreceptor debris and absorb UV light; both functions lead to high reactive oxygen species (ROS) accumulation and thus require exquisite redox balance management (Seagle et al., 2005). Cell stress resulting from ROS accumulation causes RPE dysfunction, in turn leading to visual impairment (B Domènech and Marfany 2020). Primary malfunctions of RPE can aggravate visual cell loss and lead to blindness (R. Sparrrow et al. 2010). The RPE and the neuroretina are both derived from embryonic neuroepithelial tissue and co-differentiated during the development of the optic cup (George et al. 2021). The integrity of the choroid (and retina) is partly dependent upon the generation of major RPE-derived growth factors such as IGF1, TGF-b, FGF-2, or VEGFA (Strauss 2005). Hence, the growth and function of the highly vascularized choroid is to a large extent governed by the RPE (Nickla and Wallman 2010).

In the sub-retina, composed of the RPE and the choroid, nearly 80% of glucose is converted into lactate, compared to 20% conversion in the inner retina (Wang L et al. 1997). RPE cells use lactate as fuel for ATP production via oxidative phosphorylation; this is presumed to recycle lactate and permit higher glucose availability for the photoreceptors (Viegas and Neuhauss 2021). Strikingly, metabolites such as succinate and lactate have been identified as determinants of retinal vascular development (Sapieha et al. 2008; Madaan et al. 2019). As the endogenous ligand of the G-protein-coupled receptor (GPCR) Hydroxycarboxylic acid receptor 1 (HCAR1, also known as GPR81), lactate is an important signaling molecule (Ahmed et al. 2010). More specifically, lactate-HCAR1 signaling plays a role in several physiological and pathological processes relevant to the eye, including lipid metabolism (Cai et al. 2008; Liu et al. 2009), inflammatory response (Hoque et al. 2014; Madaan et al. 2017), glucose homeostasis (Ahmed et al. 2010), angiogenesis (Madaan et al. 2019; Chaudhari et al. 2022) and cancer (Brown and Ganapathy 2020).

Based on previous findings that HCAR1 regulates inner neuro-retinal vascular development (Madaan et al. 2019), we hypothesized that ontogenic changes in the lactate receptor would also control vascular development in the sub-retina (choroid). The abundant metabolic generation of lactate in the sub-retina along with the critical role of the RPE in sustaining the choroid, led us to explore the role of the lactate receptor, HCAR1, during the choroidal development using the *Hcar1^−/−^* mouse model. We further dissected in *ex vivo* and *in vivo* experiments the underlying molecular mechanisms that are affected by HCAR1 in relation to choroidal integrity.

## Results

### HCAR1 signaling in RPE cells promotes angiogenesis in the developing choroid

Immunohistofluorescence (IHF) revealed localization of HCAR1 specifically in RPE (colocalizing with RPE65) in young pups (**Fig. 1a**). These results were corroborated by HCAR1 mRNA expression specifically on primary RPE but not on choroid of wildtype (WT) mice (**Fig. S1a-c**); mRNA expression of *Hcar1* peaked at PT9 (**Fig. 1b**) in young pups.

**Figure 1.**
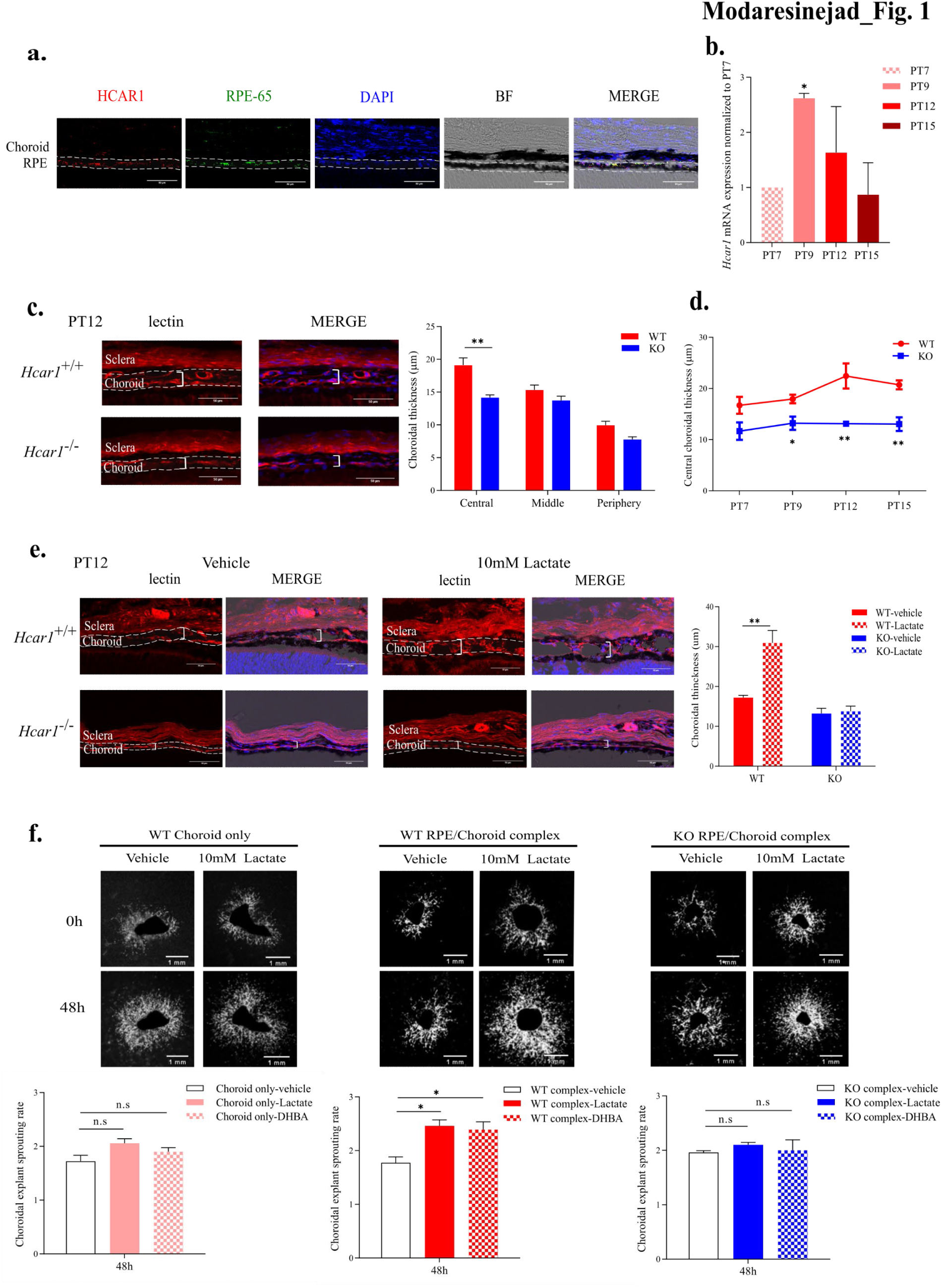
HCAR1 expression in RPE cells promotes choroidal angiogenesis (a) Representative confocal images of sub-retinas from PT12 pups, stained with anti-HCAR1 (red) and anti-RPE65 (green); nuclei were counterstained with DAPI (blue). HCAR1 and RPE signals exhibit strong colocalization. **(b)** mRNA levels of HCAR1 in the RPE/choroid layers were measured at PT7, PT9, PT12, and PT15 by RT-qPCR. **(c)** *left panel.* Representative confocal microscopy images of choroidal cross-sections from PT12 pups, stained with Texas Red-conjugated *Griffonia Simplicifolia* lectin I (red). Choroidal thickness is delineated with a dashed line; examples of measured thicknesses are shown with plain white brackets. *right panel.* Representation of the mean choroidal thickness from eyecup regions in PT12 pups. **(d)** Quantifications of central choroidal thickness at PT7, PT9, PT12, and PT15 in WT and *Hcar1*-KO (KO) mice. **(e)** *left panel.* Representative images of sub-retinas stained with Texas Red-conjugated *Griffonia Simplicifolia* lectin I (red) from PT12 mice injected intravitreally with 10 mM lactate at PT9. *right panel.* Quantification of the choroidal vasculature thickness upon lactate injection. **(f)** *upper panel.* Representative images of *ex vivo* choroid sprout assays using isolated choroid or RPE/choroid punches from WT mice, and RPE/choroid from KO mice treated with vehicle, 10 mM lactate or 80 μM DHBA for 48h. *lower panel.* Quantification of the choroid sprout assays. The explant sprouting area at 48h was normalized to the choroidal explant area at 0h. Data are presented as mean ± SEM., n=4-6 per group. *p< 0.05, **p< 0.01, Scale bar = 50 μm (a, c, e), Scale bar = 1 mm (f).

*Hcar1^−/−^* mice displayed thinner choroid at PT12 compared to their WT littermates, specifically in the central region (**Fig. 1c**). Murine choroidal thickness grows steadily post-natally to peak at 8 weeks after birth (Zhang et al. 2022). In contrast to WT mice, *Hcar1^−/−^* animals did not display any increase in critically important central choroidal thickness over the course of the first two weeks of life (**Fig. 1d**), resulting in a thinner choroid leading to substantially decreased O_2_ and nutrient delivery to the outer retina in *Hcar1^−/−^* mice (Zhou et al. 2019); growth of the peripheral choroid did not differ between WT and *Hcar1^−/−^* mice. Conversely, stimulation of the choroid of WT mice with lactate (intravitreal at PT9) triggered a robust thickening of the choroid (**Fig. 1e**), which as expected was not observed in mice deficient in HCAR1. Although the RPE harbors HCAR1, its morphology is unaltered as witnessed on phalloidin-stained flat mounts (**Fig. S1d**). On the other hand, the RPE is essential in governing the angiogenic response of the choroid to lactate stimulation. This inference was confirmed on sub-retinal explants which revealed lactate– and DHBA-(specific HCAR1 agonist (Liu et al. 2012a) induced sprouting only in explants containing both RPE and choroid, but not choroid on its own (**Fig. 1f**); RPE-choroid complex of HCAR1-null mice were unresponsive to lactate. Taken together, *in vivo* and *ex vivo* results suggest that lactate sensing by the HCAR1 receptor, residing in the RPE cells, promotes angiogenesis of immediately adjacent choroid.

### Mechanisms associated with defective choroidal growth in HCAR1-null subjects

Given that inflammation can be detrimental to sub-retina (Zhou et al. 2016; Mellal et al. 2019) and HCAR1 is reported to mitigate innate immune response (Hoque et al. 2014; Madaan et al. 2017), we looked for inflammatory cell invasion of the central sub-retina. Few Iba1+ mononuclear phagocytes were detected in both WT and HCAR1-null mice during the time when the choroid failed to grow (**Fig. S2a**). In addition, other than a rise in mRNA of pro-inflammatory *Il1b* along with that of anti-inflammatory *Il10* at PT9, other inflammatory factors were hardly changed in HCAR1-null compared to WT mice; moreover, at PT12 there tended to be a suppression of inflammatory markers in HCAR1-null animals (**Fig. S2b**). Together these observations do not support a role for inflammation in central choroidal thinning in HCAR1-null mice.

In pursuit of mechanisms potentially implicated in choroidal thinning in HCAR1-null mice, we assessed cell death. TUNEL positivity was scarcely detected in the central sub-retina (**Fig. S3a**). Likewise, local expression of the apoptotic factors *Bcl2, Bad, Casp3, Fasl,* and *Ripk3* did not differ between HCAR1-null and WT animals (**Fig. S3b**).

Since cell loss could not explain a thinner choroid in HCAR1-null mice, we surmised a slower choroidal proliferation rate, justifiably deemed in the vasculature. Concordantly, expression of the proliferation marker Ki-67 on ocular cross-sections was lower in HCAR1*-*null compared to WT mice (**Fig. 2a, S4a**). Consistently, given the critical role of RPE in eliciting choroidal vascular sprouting (**Fig. 1f**) growth factor protein expression was found to be lower in primary RPE cells from HCAR1-null mice (**Fig. S4b**). Protein microarray analysis also revealed a decrease in growth factor expression in central sub-retina of HCAR1-null vs WT mice, to some extent at PT9 and particularly at PT12 (**Fig. 2b**). Whereas, mRNA expression of a number of growth factors such as *Vegfa*, *Serpinf1*, *Sema3f*, *Fgf2*, *Angpt2* and *Igfbp2* was increased at PT9 and PT12 in HCAR1-null vs WT mice (**Fig. 2c**). We thus evaluated if depressed growth factor proteins reflected a global protein translation defect. *In situ,* sub-retinal protein translation rate determined using O-propargyl-puromycin (OPP) incorporation into proteins (Liu et al. 2012b) was substantially lower in HCAR1-null vs WT mice (**Fig. 2d, S4c**).

**Figure 2.**
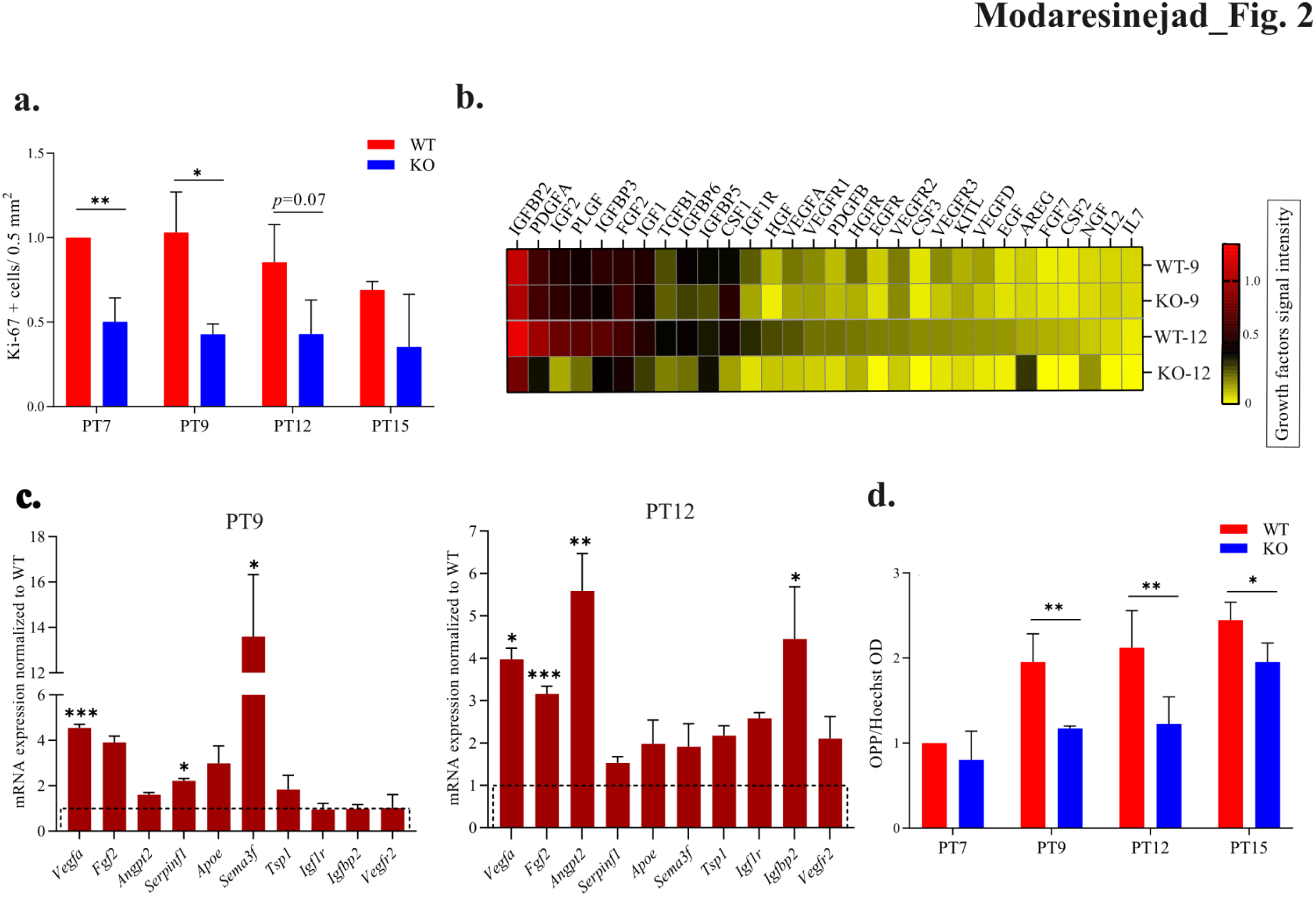
HCAR1 deficiency impairs cell proliferation and protein synthesis (a) Quantification of number of Ki-67 positive cells per 0. 5 mm^2^ in central choroid at PT7, PT9, PT12, and PT15 pups; representative images are shown in Supp Fig. 4a. **(b)** Heatmap of expression levels of several growth factors involved in cellular proliferation in WT and *Hcar1*-KO (KO) mice at PT9 and PT12. Protein levels were analyzed by protein microarray, and the signal intensity of each growth factors was normalized to the internal positive control; data was not subsequently transformed. **(c)** mRNA levels of 7 angiogenic factors were measured by RT-qPCR; data presented as fold changes of the KO relative to the WT pups, at PT9 (*left graph*) and PT12 (*right graph*). **(d)** OPP incorporation, as a proxy of protein translation rate, was revealed by the AZDye 488 Azide. 488 signal was quantified on confocal microscopy images and adjusted relatively to the intensity of the Hoechst DNA counterstain; representative images are shown in Supp Fig. 4c. Data are presented as mean ± SEM., n=3-6 per group. *p< 0.05, **p< 0.01, ***p<0.001.

Protein translation rate can be affected by various cellular stress pathways (Liu et al. 2016), notably oxidative stress (Williams et al. 2016; Quirós et al. 2017). Assessment of reactive oxygen and nitrogen species (ROS, RNS) in sub-retina of mouse pups, revealed higher levels in *Hcar1^−/−^* than in WT mice, along with the lower activity of major anti-oxidant, superoxide dismutase, and of major transcription factor regulator of redox homeostasis, NRF2 (**Fig. 3a-c**). Together these results corroborate HCAR1’s role in regulating the redox state (Akter et al. 2023) as observed herein the RPE/choroid complex during choroidal development.

**Figure 3.**
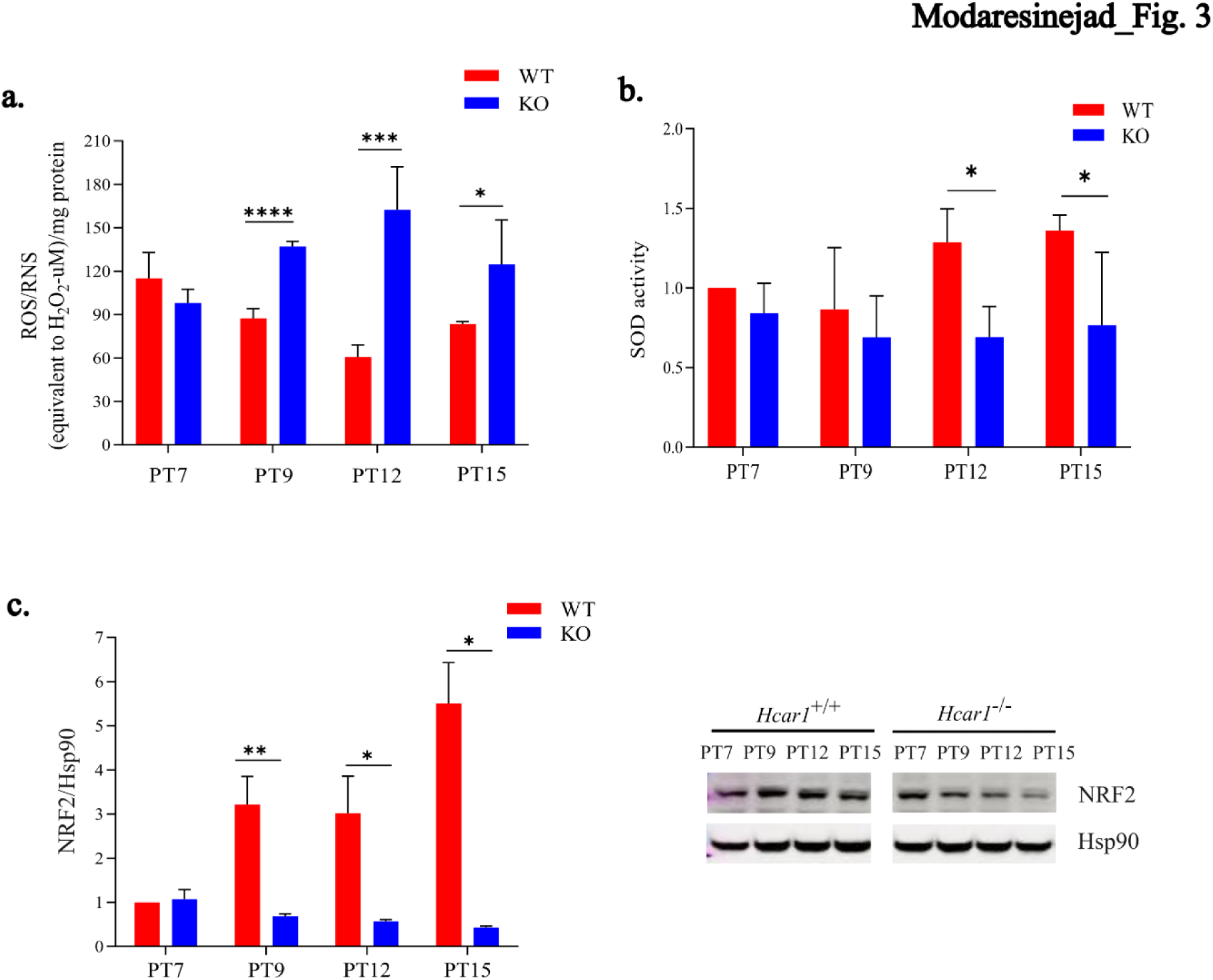
HCAR1 deficiency leads to a higher level of oxidative stress in the outer retina (a) Reactive Oxygen Species (ROS) and Reactive Nitrogen Species (RNS) were measured in freshly isolated RPE/Choroid from WT and *Hcar1*-KO (KO) pups at PT7, PT9, PT12, and PT15. ROS and RNS levels were adjusted to the amount of tissue collected. **(b)** SOD activity in freshly isolated RPE/Choroid from WT and KO pups at PT7, PT9, PT12, and PT15. **(c)** Quantification (*left panel*) and representative western blot images (*right panel*) of the stress sensor NRF2 in the RPE/Choroid from WT and KO pups at PT7, PT9, PT12, and PT15, relative to Hsp90 expression. Data are presented as mean ± SEM., n=4-6 per group. *p<0.05, **p<0.01, ***p<0.001, ****p<0.0001. Statistical analysis was performed using unpaired t-tests.

### HCAR1 deficiency and endoplasmic reticulum stress in the RPE/choroid complex

The co-occurrence of oxidative stress with choroidal involution during the second week of postnatal life led us to envisage cell dysregulation impacting proliferation. Discrepant translation and transcription (Ron 2002; Pakos-Zebrucka et al. 2016) in the phase of oxidant stress made us consider endoplasmic reticulum (ER) stress (**Fig. 4a**) (Liu et al. 2016; Cheng et al. 2016; Williams et al. 2016). Major ER stress markers BiP/GRP78 and protein disulfide isomerase (PDI) notably at the RPE (**Fig. 4b,c**), were clearly increased along with activation of cytoprotective-attempted Unfolded Protein Response (UPR) IRE1 and particularly rapidly of PERK pathways (**Fig. 4b,d,e,f**), which lead to phosphorylation and in turn inhibition of the global translation eukaryotic Initiation Factor 2α (eIF2α) (Matts and London 1984; Duncan and Hershey 1987) (**Fig. 4g**); whereas ATF6 pathway was hardly affected (**Fig. 4h**). Taken together, *Hcar1* KO leads to an increased ER stress and activation of the downstream UPR response in the sub-retina.

**Figure 4.**
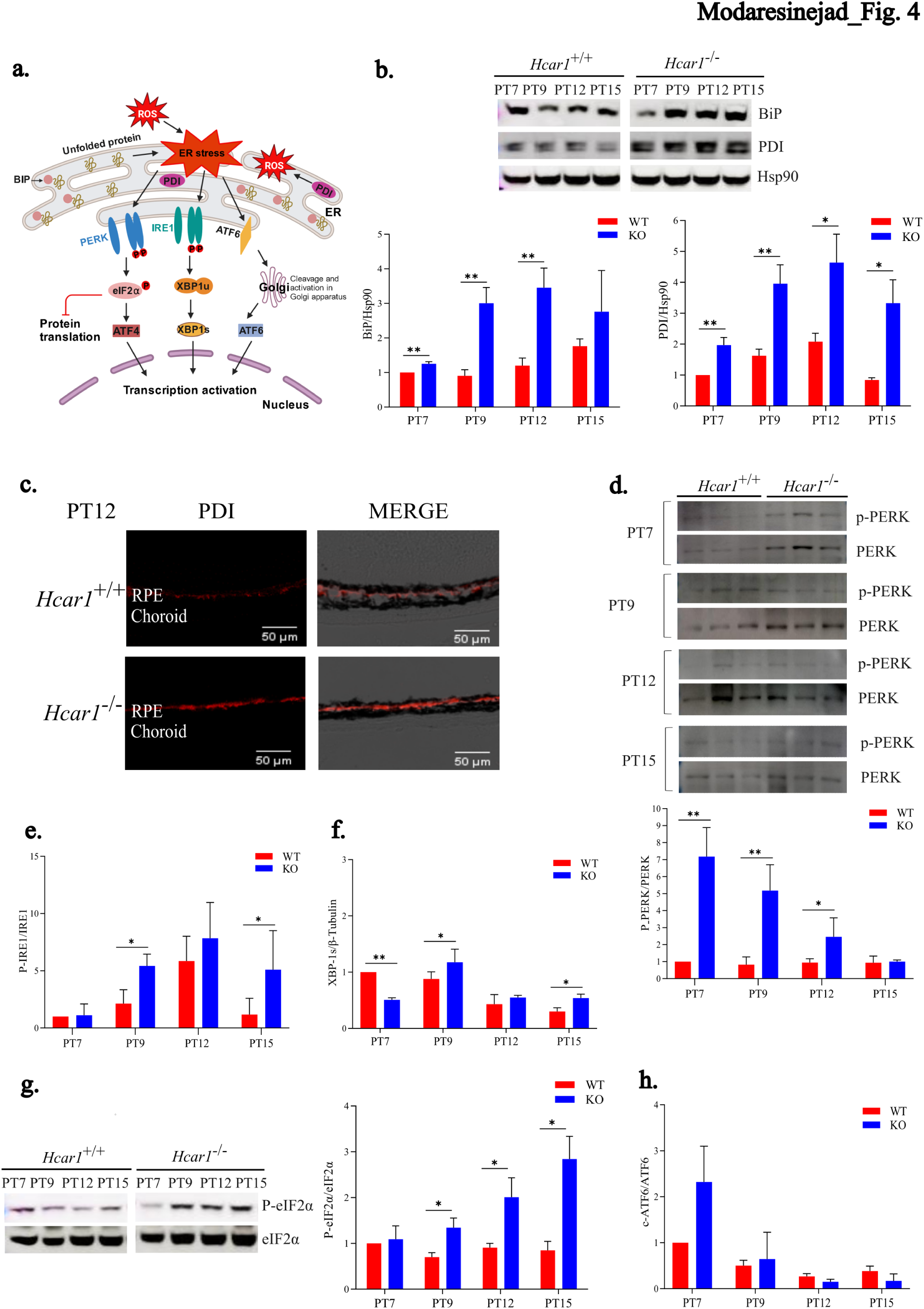
HCAR1 maintains ER homeostasis in the RPE/choroid complex (a) Schematic overview of the Unfolded Protein Response pathway. **(b)** Representative western blot images for the ER stress markers BiP and PDI, from the sub-retina of WT and *Hcar1*-KO (KO) mice at PT7, PT9, PT12, and PT15. Immediately below is quantification of the protein level for BiP and PDI. **(c)** Representative confocal images of sub-retinas from PT12 pups, stained with anti-PDI (red). **(d)** Representative western blots of phosphorylated ER stress signal PERK, from the sub-retina of WT and *Hcar1^−/−^* mice at PT7, PT9, PT12, and PT15. Immediately below is the quantification of phospho-PERK relative to total PERK. **(e)** Quantification of the phospho-IRE-1α (P-IRE-1α) relative to the total IRE-1α signal, in the sub-retina of WT and *Hcar1^−/−^* mice at PT7, PT9, PT12, and PT15. Representative images are in Supp. Fig. 5a. **(f)** Quantification of the level of spliced XBP1 (XBP-1s) relative to the level of β-Actin, in the sub-retina of WT and *Hcar1^−/−^* mice at PT7, PT9, PT12, and PT15; for representative image see Supp. Fig. 5b. **(g)** Representative western blot of phospho-eIF2α relative to total eIF2α in sub-retina of WT and *Hcar1^−/−^*mice at PT7, PT9, PT12, and PT15. Histogram is the quantification of eIF2α phosphorylation western-blot, presented as the ratio of the phospho-eIF2α signal to the total eIF2α signal. **(h)** Quantification of cleaved ATF6 (c-ATF6) relative to unprocessed ATF6, in the sub-retina of WT and *Hcar1^−/−^* mice at PT7, PT9, PT12, and PT15; representative images are in Supp. Fig. 5c. Data are presented as mean ± SEM., n=4-6 per group. *p< 0.05, **p< 0.01, ***p<0.001.

### ISR inhibitor reverses translation inhibition and choroidal involution in *Hcar1^−/−^* mice

The integrated stress response (ISR) is a complex adaptive signaling pathway that responds to diverse stresses, including extrinsic (e.g., hypoxia) and intrinsic factors (such as oxidative and ensued ER stress) (Pakos-Zebrucka et al. 2016). eIF2a is the core factor in the ISR, allowing cells to adapt to environmental and pathological stimuli (Pakos-Zebrucka et al. 2016). In addition to causing cap-dependent global inhibition of translation, eIF2a phosphorylation (Gerlitz et al. 2002) induces translation of cap-independent specific downstream effectors of the ISR, among which the transcription factor Activating Transcription Factor 4 (*Atf4*) is the best characterized (Chan et al. 2013). Western-blot analyses of ATF4 levels in the RPE/choroid of *Hcar1^−/−^* KO pups revealed increased ATF4 expression compared to age-matched WT counterparts (**Fig. 5a**). In addition, target genes downstream of ATF4 including pro-survival asparagine synthetase (*Asns*), tribbles homolog 3 (*Trib3*) and C/EBP homologous protein (*Chop/Ddit3*) were coincidentally increased at PT12 (Rozpedek et al. 2016; Gwinn et al. 2018) (**Fig. 5b**), consistent with increased activation of the ISR.

**Figure 5.**
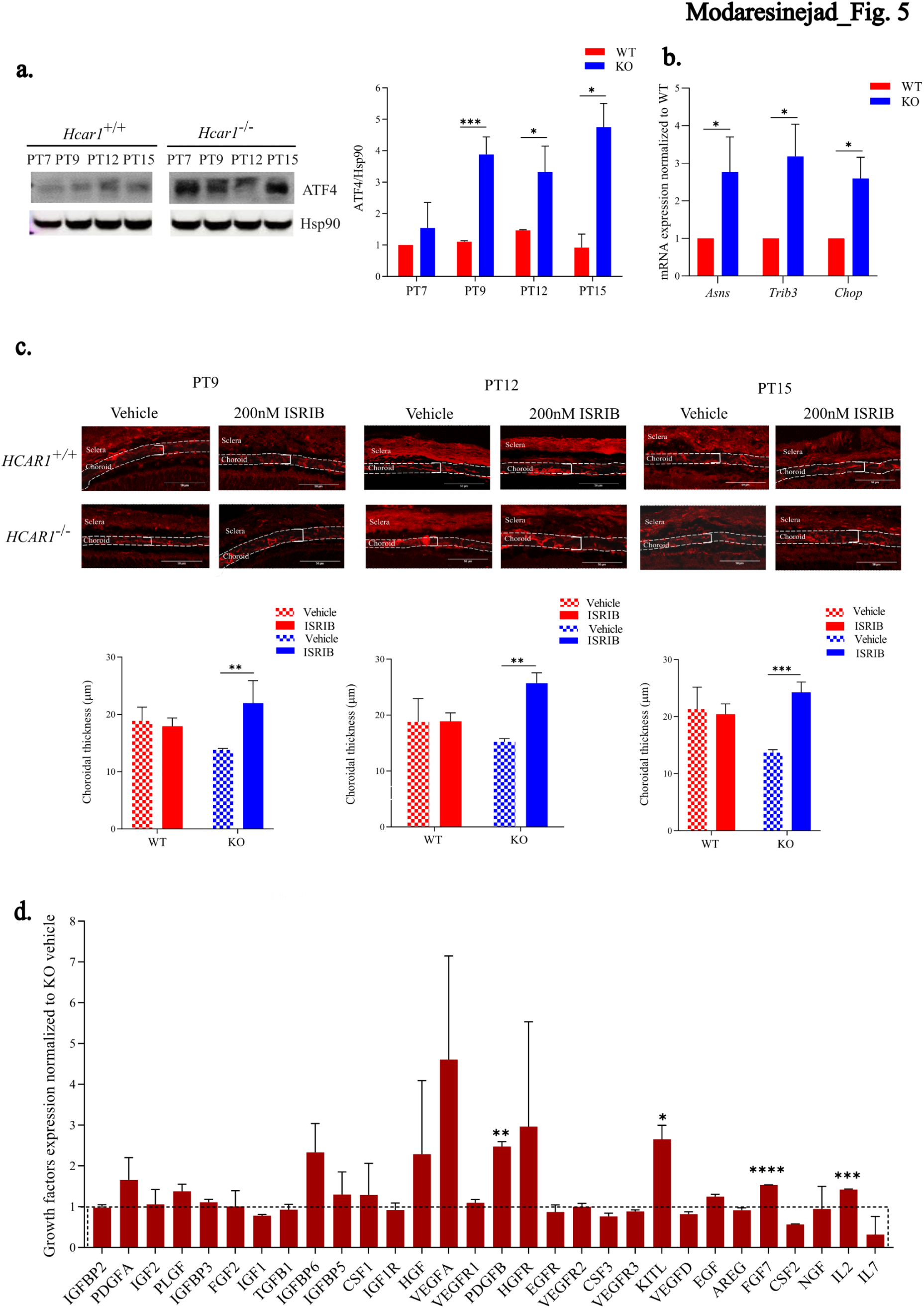
ISR inhibitor rescues the choroidal involution in *Hcar1* KO mice (a) Representative western blot and quantifications (histogram) of ATF4, from the sub-retina of WT and *Hcar1*-KO (KO) mice at PT7, PT9, PT12, and PT15. **(b)** RT-qPCR quantifications of ATF4 downstream target genes expression in the sub-retina of *Hcar1^−/−^* and WT mice at PT12. **(c)** *upper panel*. Representative confocal images of a Texas-Red-conjugated GSL-I staining of choroidal sections from PT9, PT12, or PT15 mice intravitreally-injected with ISRIB (200 nM) or vehicle at PT7, PT9, and PT12, and then sacrificed at PT9, PT12, or PT15, respectively. *lower panel*. Choroidal thickness quantifications for each corresponding age and condition. **(d)** Growth factors expression at PT12 upon injection of ISRIB at PT9 in KO mice, relative to similarly-treated WT animals. Data are presented as mean ± SEM., n=3-6 per group. *p<0.05, **p<0.01, ***p<0.001, ****p<0.0001.

We reasoned that inhibition of the ISR, which should restore protein translation and ensuing cell proliferation, would rescue choroidal thinning in the *Hcar1^−/−^* mice. For this purpose, we used the ISR inhibitor (ISRIB), a small molecule that renders the assembly of the pre-initiation complex insensitive to the phosphorylated eIF2α, thus enabling translation (Sidrauski et al. 2015; Halliday et al. 2015). ISRIB was injected intravitreally into PT7, PT9, and PT12 mice, and choroidal thickness was evaluated at PT9, PT12, and PT15. While as expected WT mice were unresponsive to ISRIB, latter led to a significant increase in the choroidal thickness in *Hcar1*-KO mice at all ages studied (**Fig. 5c**). Inhibition of the ISR pathway at PT12 in *Hcar1^−/−^* mice resulted in a rescue of the protein expression of several growth factors lost upon *Hcar1-*KO such as PDGFB, KITL, FGF7, and IL2, and normalized expression of many others (**Fig. 5d**), compared to suppressed growth factors in untreated animals (**Fig. 2b**); as expected WT mice were unresponsive to ISR inhibition. Together, observations confirm that eIF2α phosphorylation associated with ISR restricts normal choroidal development in phase of depleted *Hcar1*.

## Discussion

Integrity of the choroid is essential for vision; conversely choroidal involution has dire consequences on vision acuity as is the case in aging with macular degeneration, diabetic retinopathy, and in young subjects with retinopathy of prematurity (Nickla and Wallman 2010; Chirco et al. 2017; Zhou et al. 2019; Hamadneh et al. 2020). The choroid is primarily a vascular tissue estimated to contain the highest blood flow in the body, essential to supply O_2_ and nutrients to the highly metabolic outer retina, particularly as it applies to the macula. Accordingly, maintenance of choroidal vasculature requires a healthy RPE which is a major source of angiogenic factors (such as VEGFA, FGF2, TGFB1, IGF-1, PDGF) (Blaauwgeers et al. 1999; Strauss 2005; Saint-Geniez et al. 2006), necessary for the choroidal integrity. RPE cells are major metabolic generators of lactate. Along with a variety of other carbohydrate, protein, and lipid metabolites, lactate is not simply a metabolic end-product but a ligand for a specific GPCR, namely HCAR1. HCAR1 plays an important role in vascular development of the retina and brain (Madaan et al. 2019; Chaudhari et al. 2022). Based on the proximity and interaction of the choroid with the lactate-rich RPE, we surmised a role for HCAR1 on choroidal integrity during development. Herein, we report that HCAR1 plays a role beyond that of its ligand as a mere metabolic product – importantly as a major contributor to the development of the choroid.

HCAR1 is expressed on RPE cells (but not on choroid), yet it is essential for proper choroidal development and angiogenesis. Accordingly, stimulation of HCAR1 elicits choroidal vascular sprouting only when RPE is present adjacent to the choroid, inferring paracrine release of angiogenic factors from the RPE. Conversely, the absence of HCAR1 curtails growth factor release and in turn compromises developmental choroidal growth. The expression of HCAR1 in the RPE layer makes it an ideal location to sense the metabolic needs of the outer retina including during development and upon different light exposures (Hurley 2021). This “metabolic surveillance” mediated by the lactate receptor coordinates choroid development to permit adequate nutrient delivery to the retina; based on Poiseuille’s law the ∼35% decrease in choroidal thickness observed in HCAR1-deficient mice corresponds to a marked ∼80% drop in local hemodynamics. Hence when HCAR1 is absent, lactate sensing is impaired, leading to a decreased translation rate, thus lowering the expression of several growth factors, in turn impairing choroidal thickening. Along these lines, we previously reported that the absence of HCAR1 expression leads to curtailed inner retina vascularisation, defective transduction of photoreceptor signaling as assessed by electroretinogram, and defective retinal ganglion cell projections towards the central nervous system (Madaan et al. 2019; Laroche et al. 2021). The present study extends the concept that metabolites of different metabolic pathways have increasingly been found to exert cellular actions by acting as ligands of specific receptors as is the case for oxidative pathway intermediates (eg. HCAR1, Sucnr1, GPR99), lipid metabolism (eg. FFAR1, FFA2, FFA3, FFAR4), amino acids (eg. CaSR) and others (Mohammad Nezhady et al. 2023); HCAR1 belongs to this cluster of GPCRs.

A feature of HCAR1 deficiency downstream effects applies to curbed protein translation and ensued diminished growth factor release associated with ER stress, which is somewhat related to oxidant stress. HCAR1 signals via Erk1/2 and Akt (Mohammad Nezhady et al. 2023), which are important in growth and cytoprotection (Fujio and Walsh 1999; Tian et al. 2011) by affecting cellular energy in part via ribosome biogenesis (Shore and Albert 2022); the latter also plays a critical role in appropriate protein folding. Accordingly, defective HCAR1 expression and signaling elicit UPR. Interestingly, immunohistochemical OPP incorporation reveals that the impaired protein translation deficiency seems distributed to the RPE/choroid complex and not only the RPE, suggesting that the ER stress conveyed by the RPE impacts the growth factor-deficient choroid in HCAR1-null animals through paracrine manner. By inhibiting ER stress-triggered phosphorylated-eIF2α, critical for global protein translation, one is able to rescue developmental choroidal vascular thickness. While such rescue was coherently associated with the restoration of protein translation of various growth factors, KITL and PDGFB were notably increased; these factors exert major roles in sustaining the integrity of the vasculature (Bergers and Song 2005; Edqvist et al. 2012; Kim et al. 2019; Li et al. 2020).

The present study uncovers an unprecedented role for HCAR1 in choroidal vascular development, such that in its absence ER stress and ensuing defective protein translation interfere with the growth of this important vascular tissue. Restoration of protein translation preserves the choroid and as a consequence outer retina integrity (**Fig. S5d**).

## Material and Methods

### Animals

C57BL/6 mice were obtained from The Jackson Laboratory. *Hcar1* KO mice were purchased from Lexicon Pharmaceuticals (Texas, USA). In *Hcar1* KO mice, the transmembrane domain 2 of the coding region of *Hcar1* (100 base pairs) is replaced by a 4-kb IRES-lacZ-neo cassette (Madaan et al. 2017). Mice were maintained on standard environmentally controlled conditions (temperature: 20 ± 2 °C, humidity: 60 ± 5%, 12 h dark/ 12h light cycle). Food and water were taken *ad libitum.* All the experiments were performed in compliance with the ARVO statement for the Use of Animals in Ophthalmic and Vision Research and approved by the Animal Care Committee of the Sainte-Justine Hospital according to guidelines established by the Canadian Council on Animal Care.

ISRIB treatment was done by injection of 200 nM of ISRIB or PBS as vehicle intravitreally in PT7, PT9, and PT12 mice. Then the mice were sacrificed at PT9, PT12, and PT15, respectively and eyes were enucleated for further experiments.

### Chemical preparations

L-lactate (L7022, Sigma-Aldrich, USA) we dissolved in PBS, and the pH of the solution was adjusted with NaOH solutions (final pH 6.5-7.3; final concentration 10 mM). 3, 5-Dihydroxybenzoic acid (DHBA, D110000, Sigma-Aldrich, USA; final concentration 80 µM) was dissolved in PBS as well. ISRIB (SML0843, Sigma-Aldrich, USA) was dissolved in DMSO (final concentration 200 nM).

### RPE/choroid complex collection

Eyes were enucleated and kept in cold PBS. Then eyes were opened from the ora serrata to remove the anterior part (cornea and lens). The optic nerve was cut and the retina was separated from the posterior eye cup. For cell isolation, sterile procedures were performed. The obtained RPE/choroid complex was then dried briefly on delicate wipes followed by rapid freezing on dry ice if applicable. Eyes from a cohort of mice were prepared for experiments including RNA and protein extraction, growth factor microarray, ROS/RNS detection, and SOD activity.

### RNA extraction and Quantitative RT-PCR

Total RNA was extracted from RPE/choroid complex and primary RPE cells using a RNA extraction kit (RNeasy plus mini kit QIAGEN). The RNA concentration and integrity were determined with NanoDrop 1000 spectrophotometer. Reverse transcription was carried out using iScript cDNA SuperMix (Biorad laboratories). Primers (forward and reverse primer sequences) were designed by the Primer-Blast program and were purchased from alpha DNA or Integrated DNA Technologies (IDT). SYBR Green Master mix kit (BioRad) and Stratagene Mx3000p detection system were used to perform quantitative analysis of gene expression. qRT-PCTR assays were analyzed on triplicate samples and their threshold cycle numbers were averaged. Gene expressions were normalized to 18S expression (18S universal primer; BioRad) using the ΔCt method, and the fold change expression was analyzed by the ΔΔCt method.

### Immunohistofluorescence

Eyes or primary RPE cultures after enucleation were fixed in 4% paraformaldehyde for 1-1.5 h or 10 minutes, respectively. Eye cups were further adequately dehydrated by 30% sucrose overnight at 4°C and subsequently embedded in optimal cutting temperature (O.C.T) medium, finally, they were frozen on dry ice and kept at –80 °C. Sagittal sections of 12 μm thickness were prepared (model CM3050S cryostat; Leica Microsystems, Wetzlar, Germany) and kept at –80°C until further process. Eye cryosections or slides of RPE cells were then blocked with 1% bovine serum albumin and 0.1% TritonX-100 (T-8787; Sigma) in PBS and were afterward incubated overnight with primary antibodies followed by a secondary antibody incubation for at least 1 hour at room temperatures (Table 2). Nuclei were stained with DAPI (Invitrogen), and samples were mounted by using a gold antifade medium (P36961, ThermoFisher Scientific, USA). Slides were then analyzed by epifluorescent microscopy (E800; Nikon Eclipse, Melville, NY), or confocal microscope (SP8, Leica, Germany).

### Quantification of Choroidal Thickness

We used Griffonia simplicifolia lectin (1:500, Vector Laboratories) staining for visualizing vasculature and to designate the choroidal endothelium. Sections that contained the optic nerve head were subjected to this experiment. The choroidal vasculature thickness was measured on digital images taken by confocal microscopy under 40X amplifications using Image J software. The average of both sides around the optic nerve head was selected as the mean thickness of each sample.

### Choroidal sprout assay

We used 4-week-old mice for choroidal explants assay according to the method that was previously described (Lameynardie et al. 2005). Eye enucleations were completed under aseptic conditions and maintained in EBM-2 media (CC-3156, Lonza, USA). All connective tissue was removed from the external part of the eyes and then the cornea, lens, retina, and optic nerve head were removed from the eye cup by an incision under the ora serrata. RPE/choroid complex was further cut into 1-to 2-mm sections and placed in 30 μL Matrigel (Corning life science, USA) in a 24-well plate. After solidification of Matrigel in the incubator for 30 min, choroidal explants were kept in EGM-2 media. All these procedures were done under a laminar hood to maintain sterile conditions. Choroidal complex alone (without the RPE cell layer) was prepared based on the previous method (Defoe and Easterling 1994). Overall, the procedure for the choroidal sprout assay was like the preparation of the RPE/choroid complex except for the digestion step. After removing connective tissue, fat tissue, and blood vessels at the external part of the eye cup, the eyes were incubated with 2% Dispase II (neutral protease; Roche Applied Science, Indianapolis, IN) in Hank’s balanced salt solution (HBSS, Invitrogen) at 37 °C for 30 min. The explants were kept in the media for 3 days without any treatments until the vessel grew. On the 4^th^ day, explants were treated with PBS (vehicle), 10 mM lactate, or 80 μM DHBA until the 6^th^ day. Photos were taken prior to the treatment daily until the 6^th^ day. By using ImageJ, the angiogenesis response was determined by measuring the vessel area and normalizing to the control groups.

### Western blot analysis

Collected RPE/choroid complex from aged match tissue samples (PT7, PT9, PT12, and PT15) and from WT and *Hcar1* KO mice were lysed in RIPA buffer (cell signaling) supplemented with 0.1 mg/ml PMSF and proteinase inhibitor cocktail (11697498001, Roche, USA), and then were homogenized thoroughly (BRANSON Sonifier 150) and kept at 4°C for 1 h. After centrifuging at 12,000xg at 4°C for 15 min, the supernatant was collected and subjected to protein concentration measurement by Bradford’s method (Bio-Rad) (Bradford 1976). Around 40 μg of protein sample was loaded and electrophoresed on SDS-PAGE gel and electroblotted onto PVDF membranes. After blocking with 5% BSA, the membranes were immunoblotted overnight with primary antibodies (table 2) at 4°C. After washing, membranes were incubated with their respective secondary antibodies horseradish peroxidase (HRP)-conjugated. Enhanced chemiluminescence (GE Healthcare) was detected by using the ImageQuant LAS-500 (GE Healthcare, Little Chalfont, United Kingdom). Image analysis was done with ImageJ software.

### Growth factor microarray

We used a commercial kit to detect the level of various growth factors which are involved in angiogenesis (mouse growth factor array C3, RayBiotech, USA). After blocking the membrane at room temperature, 500 μg of total protein was loaded and incubated overnight at 4°C. After washing the membrane, a biotinylated antibody cocktail was added into each well and incubated overnight at 4°C and detected by HRP-streptavidin and chemiluminescence. Membranes were imaged using the ImageQuant LAS-500. The analysis was done with ImageJ.

### ROS/RNS detection

Levels of reactive species were determined with a commercial kit that uses an oxidizable fluorogenic probe, DCFH-DiOxyQ (STA-347-5, Cell Biolabs, USA). RPE/choroid complex was homogenized, and then 80 μg of lysate was added to a 96-well plate with a black wall and transparent bottom. The Fluorescence was read at 480 nm Ex/530 nm Em. Samples were run in quadruplicate. H_2_O_2_ dilutions were used for the standard curve. The levels of ROS/RNS were calculated and normalized to the amount of protein.

### Superoxide dismutase (SOD) detection

SOD activity was measured by a detection kit that uses the water-soluble tetrazolium salt, WST-1 [2-(4-Iodophenyl)-3-(4-nitrophenyl)-5-(2,4-disulfophenyl)-2Htetrazolium, monosodium salt] (19160, Sigma-Aldrich, USA). RPE/choroidal complex of a cohort of mice was obtained and homogenized by sonication for 10 seconds in ice-cold buffer 0.1 M Tris/HCl (pH 7.4) containing 5 mM β-ME, 0.5% Triton X-100 and 0.1 mg/ml PMSF with a tissue-weight/buffer of the ratio of 1:8 at 4 °C. After centrifuging at 14,000x g for 5 min at 4°C. Then supernatants were collected to assay according to the manual. Absorbance was read out at 450 nm. SOD activity was then calculated based on the formula provided by the kit.

### Protein synthesis rate measurement

The nascent protein synthesis rate was measured using Click-&-Go® Plus 488 OPP Protein Synthesis Assay Kit (Click-chemistry tool) according to the manufacturer’s protocol. Equal sections of the sub-retina were dissected and washed in a 6-well plate with media. After 5 hours OPP was added which is an analog of puromycin (can enter the acceptor site of ribosomes and incorporates into nascent polypeptide chains) and then samples were incubated for 30 min. The amount of incorporated OPP is identified by a click chemical reaction by AZDye 488 Azide (Alexa Fluor 488® equivalent). Finally, the intensity of AZDye 488 Azide is adjusted with the intensity of DNA counterstain Hoechst 33342 and indicates the nascent protein synthesis rate.

### Statistical analysis

Statistical analysis was carried out by Prism software (GraphPad Software). and two-tailed unpaired Student’s t-test and one-way ANOVA were used for data analysis. Significance between groups was calculated with Bonferroni post hoc analysis. Data were presented as means ± SEM with the statistical significance at *P<0.05, **P<0.01, ***p<0.001, ****p<0.0001.

## Acknowledgment

This work was supported by the Canadian Institutes of Health Research grant. M. M. was supported by scholarships from the faculty of Medicine Université de Montréal. X. Y. was supported by scholarships from the China Scholarship Council and the School of Optometry at the University of Montreal. M. A. M. N. was supported by S.Véronneau-Troutman and Université de Montréal Ophthalmology department, program in molecular biology and faculty of medicine scholarships. We thank Dr. E. Kuster from the microscopy facility of CR Sainte Justine Hospital and Dr. X. Hou for their input on the work presented here. Also, we thank Sonja Lesperance for her input in the animal facility. S. C. holds a Canada Research Chair (Translational Research in Vision) and Leopoldine Wolfe Chair Vision.

## Author Contributions

M. M. and X. Y. designed the study, performed and analyzed the experiments, and drafted the manuscript equally. M. A.M. N. designed the study, performed the analysis of the data, and drafted the manuscript. X. H. helped with performing the animal study. E. B. and J. C. R. analyzed data and E.B. was involved in writing the manuscript. T.Z. and H.T. performed experiments. P. L., P. H., and S. C. conceived and supervised the whole project and oversaw the conception, experiments, analysis, and writing of the work.

